# Endogenous *Staphylococcus aureus* CRISPR-*cas* system limits phage proliferation and efficiently excises from the genome as part of the SCC*mec* cassette

**DOI:** 10.1101/2023.03.19.533347

**Authors:** Kasper Mikkelsen, Janine Zara Bowring, Yong Kai Ng, Frida Svanberg Frisinger, Julie Kjærsgaard Maglegaard, Qiuchun Li, Raphael N. Sieber, Andreas Petersen, Paal Skytt Andersen, Jakob T. Rostøl, Nina Molin Høyland-Kroghsbo, Hanne Ingmer

**Affiliations:** Department of Veterinary and Animal Sciences, University of Copenhagen, Denmark; Department of Bacteria, Parasites and Fungi, Statens Serum Institut, Copenhagen, Denmark; Jiangsu Key Lab of Zoonosis/Jiangsu Co-Innovation Center for Prevention and Control of Important Animal Infectious Diseases and Zoonoses, Yangzhou University, China; Centre for Bacterial Resistance Biology, Imperial College London, London SW7 2AZ, UK; Department of Plant and Environmental Sciences, University of Copenhagen, Denmark

**Keywords:** CRISPR-Cas Type III-A, *Staphylococcus aureus*, MRSA, SCCmec type V(5C2&5), bacteriophage

## Abstract

CRISPR-Cas is an adaptive immune system that allows bacteria to inactivate mobile genetic elements. Approximately 50% of bacteria harbor CRISPR-*cas*, however in the human pathogen *Staphylococcus aureus*, CRISPR-*cas* loci are less common and often studied in heterologous systems. We analyzed the prevalence of CRISPR-*cas* in genomes of methicillin resistant *Staphylococcus aureus* (MRSA) isolated in Denmark. Only 2.9 % of the strains carried CRISPR-*cas* systems, but for strains of sequence type ST630 over half were positive. All CRISPR-*cas* loci were type III-A and located within the staphylococcal chromosomal cassette (SCC*mec*) type V(5C2&5) conferring β-lactam resistance. Curiously, only 23 different CRISPR spacers were identified in 69 CRISPR-positive strains and almost identical SCC*mec* cassettes, CRISPR arrays and *cas* genes, are present in staphylococcal species other than *aureus*, suggesting that these were transferred horizontally. For the ST630 strain 110900, we demonstrate that the SCC*mec* cassette containing CRISPR-*cas* excises from the chromosome at high frequency. However, the cassette was not transferable under the conditions investigated. One of the CRISPR spacers targets a late gene in the lytic bacteriophage (phage) virus philPLA-RODI, and we show that the system protects against phage infection by reducing phage burst size. However, CRISPR-Cas can be overloaded or bypassed by CRISPR escape mutants. Our results imply that the endogenous type III-A CRISPR-Cas system in *S. aureus* is active against targeted phages, albeit with low efficacy. This suggests native *S. aureus* CRISPR-Cas offers only partial immunity, and in nature may work in tandem with other defense systems.

**Importance:** CRISPR-Cas is an adaptive immune system enabling bacteria and archaea to protect themselves against mobile genetic elements such as phages. In strains of *Staphylococcus aureus*, CRISPR-*cas* is rare, but when present, it is located within the SCC*mec* element encoding resistance to methicillin and other β-lactam antibiotics. We show that the entire module is excisable, with almost identical versions found in different species of *non-aureus* staphylococci suggesting that the system only rarely acquires new spacers in *S. aureus*. Additionally, we show that in its endogenous form, the *S. aureus* CRISPR-Cas is active but inefficient against lytic phages, with phages being able to form escape mutants or overload the system. This leads us to propose that CRISPR-Cas in *S. aureus* offers only partial immunity in native systems, and so may work together with other defense systems to prevent phage-mediated killing.

## Introduction

Clustered regularly interspaced short palindromic repeats (CRISPR) and their CRISPR associated Cas proteins are microbial adaptive immune systems that protect against invading genetic elements such as phages [1] and plasmids [2]. Genetic memory of prior encounters is stored as spacer sequences in the CRISPR array and adaptation is accomplished by acquisition of foreign DNA fragments that are inserted as new spacers downstream of the CRISPR leader region [3]. The CRISPR array is transcribed as a pre-CRISPR RNA that is processed by Cas proteins into the mature CRISPR RNAs (crRNAs). During the interference stage, the crRNAs guide Cas protein complexes to the foreign DNA or RNA sequences by binding matching protospacers, with subsequent invader nucleic acids being destroyed by the Cas nucleases [4].

Despite its ability to protect bacteria from foreign genetic elements, only approximately 50% of all bacteria harbor CRISPR-Cas immune systems [5], and for some bacterial species the system is only present in a subset of strains. One such example is *S. aureus*. It is an opportunistic, human pathogen that naturally colonizes both humans and animals and gives rise to serious, life-threatening infections. Until now, only a few *S. aureus* strains have been reported to carry CRISPR-*cas* systems and they almost exclusively belong to the type III-A sub-group that targets both DNA and RNA [6–9]. The type III-A CRISPR-Cas activity is dependent on transcription of target sequences, and involves specific RNase activity recognizing transcripts of these sequences as well as single-stranded DNase activity, degrading the target DNA [10, 11]. Much of the previous work characterizing the type III-A CRISPR-Cas systems in *S. aureus* has been performed using expression vectors or heterologous expression systems [10, 12, 13] and thus, little is known about the activity of the endogenous *S. aureus* type III-A CRISPR-Cas systems against invading elements [7].

In *S. aureus*, strains are subdivided based on multi-locus sequence types (STs) and on composition of the *spa* gene, and they are categorized as being either hospital-, community-or livestock-associated [14, 15]. The first CRISPR-Cas element described in *S. aureus* was reported for a livestock-associated, MRSA strain 08BA02176, with the sequence type ST398 and *spa* type t034 [6]. 08BA02176 was isolated from a human infection in Canada, and it carried an intact CRISPR-*cas* locus with 15 spacers located within the CRISPR array upstream of the *cas* genes and 3 spacers in the downstream array. Interestingly, the CRISPR-*cas* element was located within the SCC*mec* cassette that in MRSA strains carries the *mecA* gene encoding the alternative penicillin-binding protein providing methicillin resistance. In another study, five clinical isolates of MRSA were demonstrated to carry a type III-A CRISPR-*cas* system located in the SCC*mec* type V cassette [8]. This study showed that type III-A CRISPR-Cas activity depends on transcription and that Cas10, Csm2, Csm3, Csm4 and Cas6 are required for CRISPR-Cas interference. We recently examined available *S. aureus* genome sequences for CRISPR-*cas* systems and found 35 genomes to encode a complete type III-A CRISPR-Cas system [7]. In all cases the system was located within the SCC*mec* cassette and for one strain we tested the CRISPR-Cas activity and demonstrated CRISPR-Cas dependent protection against phage infection.

In addition to *S. aureus*, CRISPR-*cas* systems also appear in other staphylococcal species including *Staphylococcus argenteus*, a species recently re-classified from *S. aureus* strains belonging to ST1850 and ST2250 [8, 16, 17]. In the coagulase negative staphylococci (CoNS), type III-A systems are found in *S. epidermidis, S. lugdunensis, S. capitis* and *S. warneri* [18, 19]. Curiously there is extensive homology of *cas* genes and CRISPR spacers between *S. aureus* and CoNS leading to the suggestion that there may have been recent exchange of CRISPR-*cas* along with SCC*mec* between the bacterial species [9, 20].

Here, we report the prevalence of CRISPR-*cas* systems in clinical MRSA strains in Denmark, and show that type III-A CRISPR-*cas* systems are present in more than half of the examined strains belonging to the emerging clone, ST630 [21, 22]. As in previous studies, the CRISPR-*cas* system is located within the SCC*mec* cassette type V(5C2&5) and has substantial homology to similar elements in the CoNS. Interestingly, we demonstrate that the entire SCC*mec* cassette excises and circularizes at high frequencies, suggesting that the circular form can be transferred horizontally and that recipient strains acquire CRISPR-*cas* phage defense and methicillin resistance simultaneously. Yet, we did not detect transmission of the SCC*mec* cassette between *S. aureus* strains. Further we show that the endogenous CRISPR-cas system in the clinical isolate 110900 protects against the lytic phage philPLA-RODI, but it can be circumvented either by escape mutations in the phage, or when the system is overloaded with phage. This implies that the endogenous type III-A CRISPR-cas systems of *S. aureus* offer varying degrees of protection against phage infections, even when encoding a CRISPR spacer targeting the phage.

## Results

### CRISPR-*cas* prevalence in Danish clinical *S. aureus* isolates

To determine the prevalence of CRISPR-*cas* systems in Danish clinical *S. aureus* isolates, we analyzed 1504 clinical MRSA isolates sequenced at Statens Serum Institut (SSI). We screened the strains for the presence of the conserved *cas1* and *cas2* genes [23] and typed the CRISPR systems using the CRISPRCasFinder database [24]. Of the initial 1504 isolates, 54 (3.6%) contained a complete CRISPR-*cas* system, defined as a full set of *cas* genes flanked by two CRISPR arrays, all of which belong to the type III-A system. The highest prevalence was found in ST types 2250 and 1850, but as these were recently re-annotated as *S. argenteus* [17] they were excluded from the analysis, as was one ST5 isolate that proved to be *S. epidermidis*. Thus, the resulting frequency of MRSA CRISPR-positive (CRISPR^+^) isolates was reduced to 2.9% (43/1490). The most prevalent CRISPR^+^ *S. aureus* clone is ST630 with more than half of the isolates carrying CRISPR-*cas*, followed by ST1, ST5 and ST130 (Table 1). As ST630 is an emerging clone in both Denmark and China [21, 22, 25], we decided to include 30 additional ST630 isolates sequenced after 12^th^ of March 2019. We furthermore excluded a few isolates due to inadequate CRISPR array sequencing, bringing the total number of CRISPR^+^ strains to 69.

**Table 1.** CRISPR-*cas* prevalence in clinical *S. aureus* isolates. Number of CRISPR^+^ *S. aureus* isolates investigated and their distribution across ST types.

| ST Type | CRISPR<sup>+</sup> strains/Total | CRISPR<sup>+</sup> strains (%) |
| --- | --- | --- |

|  | number of strains |  |
| --- | --- | --- |
| 1 | 13/183 | 7.1 |
| 5 | 5/132 | 3.8 |
| 130 | 2/83 | 2.4 |
| 630 | 22/40 | 55 |
| 3208 | 1/1 | 100 |

Interestingly, for all the *S. aureus* CRISPR^+^ clinical isolates, the type III-A CRISPR systems are located in *SCC*mec** cassettes of the type V(5C2&5) around 3-5 kb downstream of the *ccrC1-* allele-2, within the SCC*mec* joining region 1 (J1 region), which is a preferred integration site for plasmids and transposons carrying antibiotic resistance genes [26, 27]. In addition to CRISPR-*cas*, the J1 region contains a transposon with the enterotoxin H gene followed by a sulfite/sulfate resistance cassette in ST1 strains, the ST5 strains carry a sulfite/sulfate resistance cassette, and a *kdp* operon and an arsenite/arsenate resistance cassette is present in ST630 strains. Thus, CRISPR-*cas* loci in *S. aureus* are generally found inside the J1 region of the type V(5C2&5) *SCC*mec** cassette from isolates of various clonal complexes [7, 25].

### Spacer patterns of the *S. aureus* type III-A CRISPR arrays

Among the 69 Danish clinical CRISPR^+^ isolates and the one reference strain (ST398 isolate 08BA02176) we found 23 distinct spacers present in 37 CRISPR arrays with 20 spacers in CRISPR array 1 (spacers 1.1-1.20) and three in CRISPR array 2 (spacers 2.1-2.3) (Figure 1).

**Figure 1.**
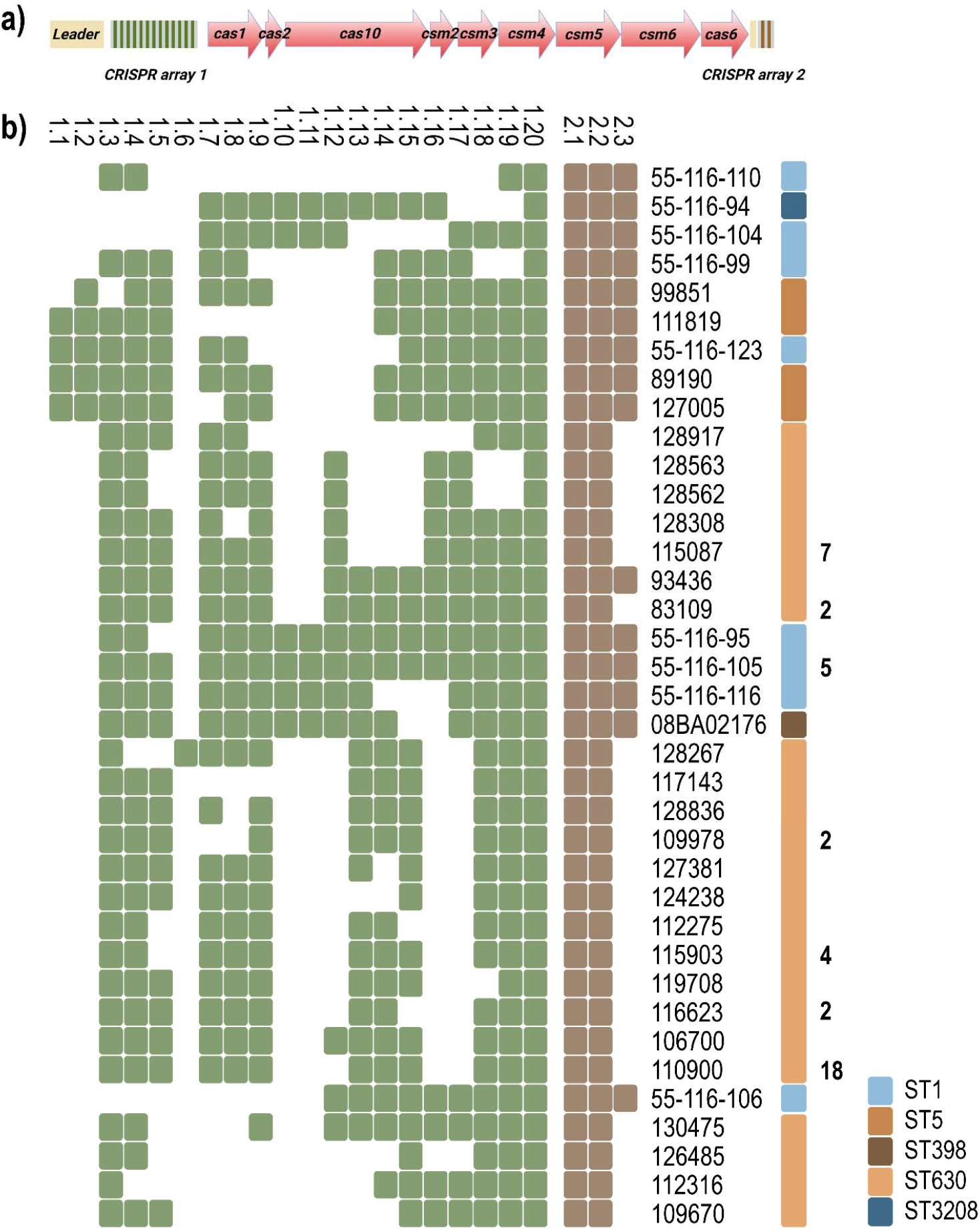
Overview of the type III-A CRISPR-*cas* systems examined in *S. aureus*, **(a)** Graphic representation of the type III-A CRISPR-*cas* system including leader sequences (yellow), *cas* and *csm* genes (red) and CRISPR array 1 (green/grey) and 2 (brown/grey). **(b)** Spacer content in CRISPR array 1 and 2 in 69 Danish clinical isolates and 1 reference strain (08BA02176). Green and brown squares symbolize the presence of the spacer in CRISPR array 1 and 2, respectively. STs are noted by color and the number of isolates that carry identical CRISPR arrays is indicated in bold where no number indicates 1 strain. The order of the spacers is presented as they occur in the array.

In active CRISPR-Cas systems, spacer acquisition commonly occurs at the 5’ end of the CRISPR array, immediately downstream of the CRISPR leader sequence. This results in highest diversity at this end of the array when comparing CRISPR arrays with a recent common origin [28, 29]. Five of our isolates had spacers 1.1 and 1.2 as 5’ spacers, which suggests recent incorporation of new spacers in these arrays, as 62 of 70 isolates we analyzed had spacer 1.3 as 5’ spacer and was succeeded by spacer 1.4 in 60 of these arrays (Figure 1b). The remaining three isolates had middle array spacers from other isolates as their 5’ spacers, suggesting that these had not adapted since diverging from the common array.

The 3’ end of CRISPR arrays is commonly conserved among strains since these spacers have resided there the longest, until they are eventually lost [30]. Accordingly, we found the 3’ end to be highly conserved with all isolates carrying spacer 1.20 as the final 3’ spacer and 64 of the 70 isolates carrying spacers 1.18 and 1.19 immediately upstream. Surprisingly, we observed the highest diversity in the center of the array, ranging from isolate 55-116-110 that only hold 5’- and 3’-end spacers of other strains in its CRISPR 1 array, to others (*e.g*. 55-116-105) containing nearly all middle-array-spacers (Figure 1).

Next, we compared CRISPR spacer composition with ST clonality. We found that CRISPR arrays in ST630 isolates have the 1.3 spacer at its 5’ end, while ST5 arrays start with either the 1.1 or 1.2 spacers. Additionally, the 1.10 and 1.11 spacers are limited to ST1, ST3208 and ST398 isolates. Similarly, for the CRISPR array 2, all but one ST630 isolate carry only the 2.1 and 2.2 spacers and all non-ST630 strains carry an additional 2.3 spacer. These observations along with the lack of diversity in the 5’ end of CRISPR array 1, and the presence of only 23 different spacers in the two CRISPR arrays shared between 70 strains, suggest that the arrays share common origin via horizontal transfer and that adaptation of new spacers is infrequent and can occur at non-canonical sites e.g. in the middle of the array in these *S. aureus* strains.

### Spacers found in Danish MRSA isolates target staphylococcal phages and plasmids

To investigate the target protospacers of the CRISPR-*cas* systems in Danish MRSA isolates we searched the NCBI database for spacer homology. Here we found that 7 of the 23 spacers had homology to known mobile genetic elements (Table 2) where spacers 1.9, 1.14, 1.18, and 2.2 target phages and 1.5, 1.19, and 1.20 target plasmids. Importantly, of these 7 protospacers, 6 are located within annotated open reading frames are complementary to the respective mRNA transcripts. This reflects that type III-A CRISPR-Cas systems target transcripts of invading mobile elements [8, 31]. Spacer 1.20 covers a bidirectional promoter region but might only be functional during transcription of the *parA* gene due to the aforementioned strand bias.

**Table 2.**
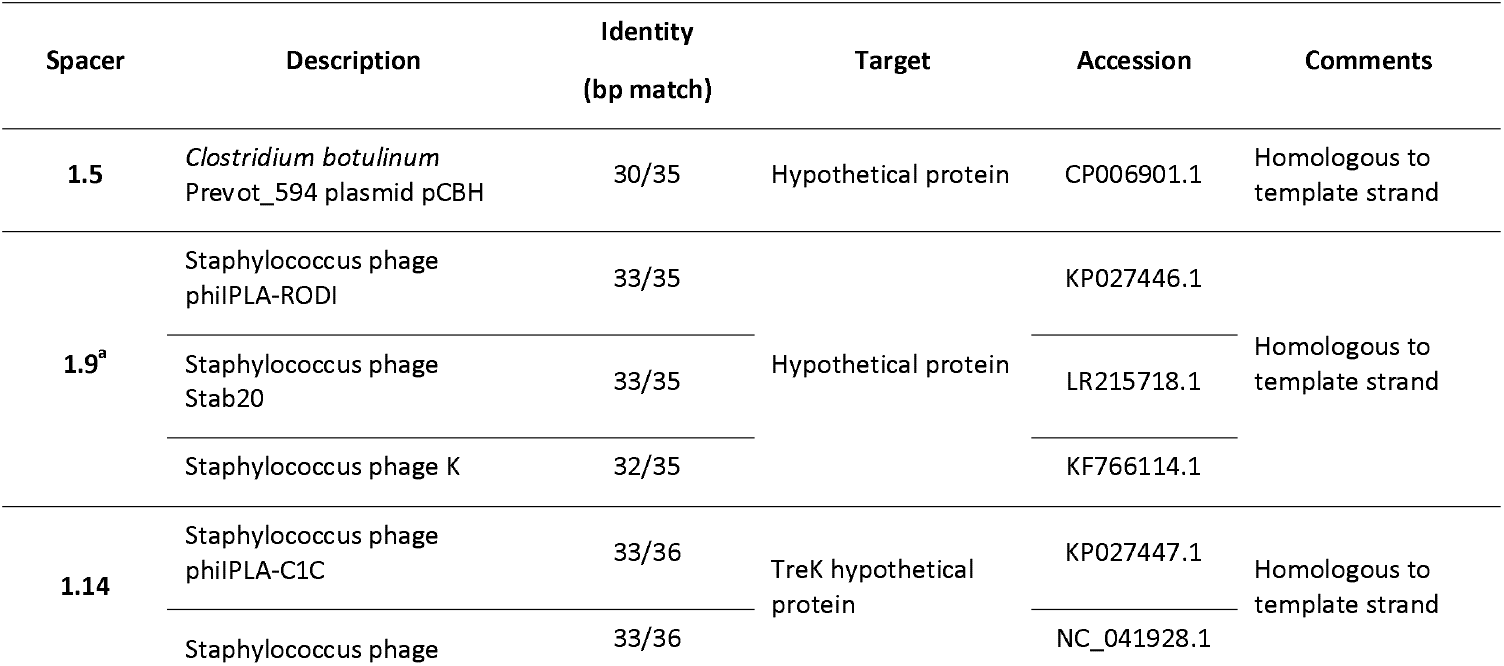
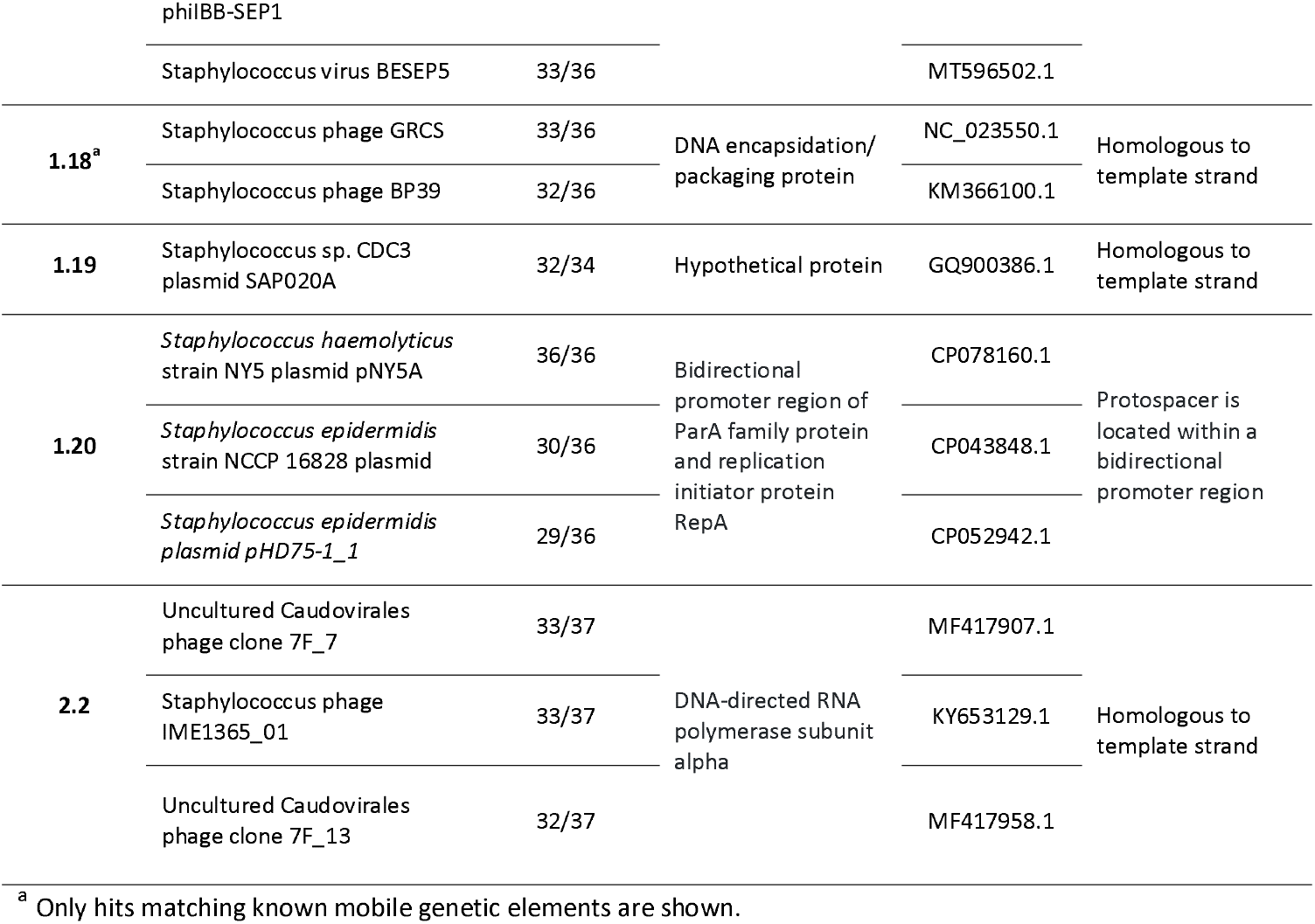
Spacer targets. Hits from a blastn search of spacers 1.1 to 2.3 from CRISPR arrays 1 and 2 against the NCBI database are shown with annotated identity, and protein and strand targets.

The spacer length ranges from 32 to 39 bp and often carry two or more mismatches to the identified protospacer sequence (Table 2). However, complete viral escape from the type III-A system was shown only to arise through complete target deletion [32], why we would assume these spacers to carry at least some degree of activity.

### Type III-A CRISPR-*cas* is located in the SCC*mec* cassette and is present in various staphylococcal species

To further explore type III-A CRISPR-*cas* homology within staphyloccoci, we blasted the sequence covering *cas1-6* of the *S. aureus* strain 110900 [22] against the NCBI database. Based on the hits, we constructed a phylogenetic tree and compared them based on their core genomes (Figure 2a). Also, we assessed their *cas* gene identity (% identity compared to the *cas1-6* sequence of 110900), *SCC*mec** subtypes, and whether they carried any homologues of the 23 *S. aureus* spacers identified in our screen (1.1 to 2.3). We found that *S. aureus* strains share almost identical *cas* gene sequences, with >99.9 % identity at nucleotide level. Interestingly, other staphylococcal species as diverse as *S. capitis, S. schleiferi*, and *S. pseudintermedius* likewise contain nearly identical *cas* gene sequences (>99.9 %) to the *S. aureus* 110900 strain. Additionally, strains of *S. argenteus* and *S. equorum* share 94% identity to the *S. aureus cas1-6* gene sequence followed by the *S. epidermidis* (89%) and *S. lugdunensis* (76%) strains (Figure 2a, *cas* similarity column).

**Figure 2.**
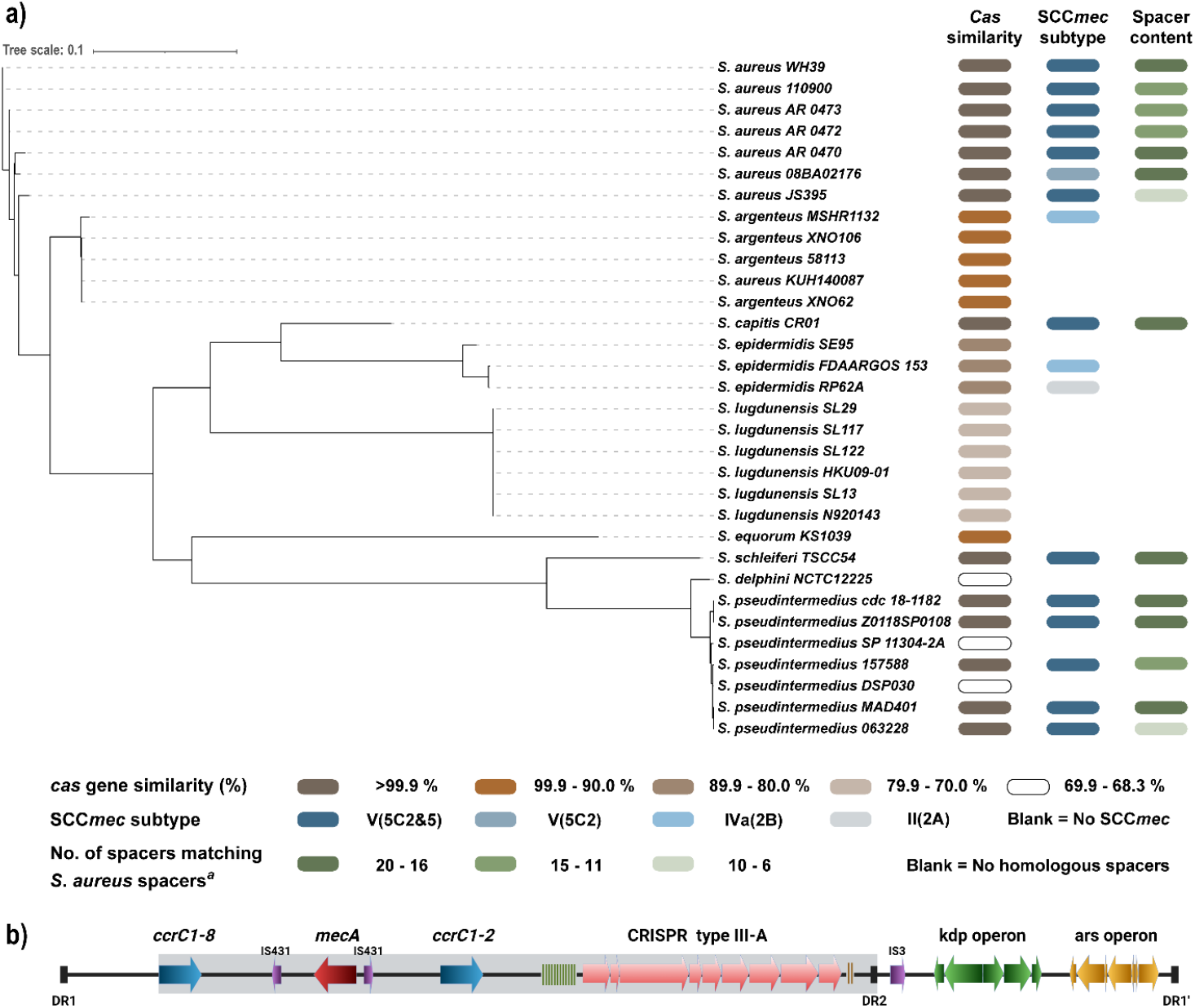
The SCC*mec* type V(5C2&5), harboring type III-A CRISPR-*cas*, is shared between staphylococcal species. (a) Phylogenetic core genome single nucleotide polymorphism (SNP) tree based on hits from a blastn search of the *cas* gene sequence (*cas1* to *cas6*) of strain 110900. *cas* gene identity (%), SCC subtype, and number of *S. aureus* spacers are indicated at the bottom. ^a)^ Sum of spacers homologous to the *S. aureus* spacers (1.1 to 2.3) found in staphylococcal isolates. (b) Graphical presentation of the SCC*mec* type V(5C2&5) with the highly conserved region marked in grey.

In addition to the *cas* locus, strains of *S. capitis, S. schleiferi*, and *S. pseudintermedius* carry *SCC*mec** type V(5C2&5) cassettes nearly identical to that of 110900 (Figure 2b). They share 99% identity in the 32 kb region from the *ccrC1-8* to Direct Repeat 2 (DR2), which includes both recombinase genes, *mecA* and the entire CRISPR-*cas* system (Figure 2b, grey shaded area). Also we find that these strains carry high numbers of homologous spacers to those found in the *S. aureus* strains screened earlier in this study (>16), including identical homologues of all of the spacers identified in the *S. aureus* strain 110900. To sum up, the high degree of identity between *S. aureus* 110900 *SCC*mec** type V(5C2&5) (including *cas* genes and CRISPR arrays) and *SCC*mec** type V(5C2&5) of other staphylococcal species strongly points to incidences of intra- and inter-species horizontal transfer of the *SCC*mec** cassette together with the CRISPR-Cas system.

Based on the core genome sequences, *S. aureus* and *S. pseudintermedius* are most distantly related (Figure 2a). Interestingly, *S. pseudintermedius* either share type III-A *cas* genes with high (>99.9%) or low (<70%) identity to the *S. aureus* strain 110900 coinciding with the CRISPR system being located in the *SCC*mec** or elsewhere on the chromosome of *S. pseudintermedius*, respectively. This further supports that the type III-A CRISPR-*cas* system found in *S. aureus* is transferable between staphylococcal species, and likely is mobilized via the *SCC*mec** type V(5C2&5).

### The SCC*mec* type V(5C2&5) cassette, including the CRISPR-*cas* locus, circularizes

*SCC*mec** cassettes in *S. aureus* have previously been shown to excise from the genome [33] and therefore, we examined if the *SCC*mec** cassette containing the CRISPR-*cas* locus was excisable in strain 110900. The *SCC*mec** is flanked by an upstream direct repeat (DR) and two downstream DRs (DR2 and DR1’, Figure 3a) that can potentially recombine to form extrachromosomal circles [34]. If so, two distinct circularized fragments may form, both of which contain the CRISPR-*cas* system along with the *mecA* gene, namely a 38 kb fragment arising via recombination of the DR1/DR2 repeats, or a 59 kb fragment via recombination of the DR1/DR1’ pair (Figure 3a). We designed primers to span the junction of the two putative circular *SCC*mec** entities and used qPCR to quantify levels of excised, circularized SCC*mec* normalized to the amount of *adsA* gene located 1kb upstream of SCC*mec*. The 59 kb fragment generated by excision of the entire SCC*mec* cassette involving DR1/DR1’ recombination showed a relatively high excision and circularization frequency of 10^-1^ relative to the abundance of the chromosomal control (Figure 3b). Next we hypothesized that specific environmental signals such as antibiotics may trigger or enhance excision, as previously observed for the SCC*mec* type V(5C2&5) in *Staphylococcus haemolyticus* [34]. However, circularization frequency was not affected by subinhibitory concentrations of the β-lactam antibiotic, oxacillin nor by the DNA damaging agent mitomycin C. The excision frequency of the 38 kb fragment was approximately 10^-7^ with or without antibiotics making it six orders of magnitude less frequent than excision of the entire cassette. This difference is probably due to the three-nucleotide mismatch between DR1 and DR2 (Figure 3c).

**Figure 3.**
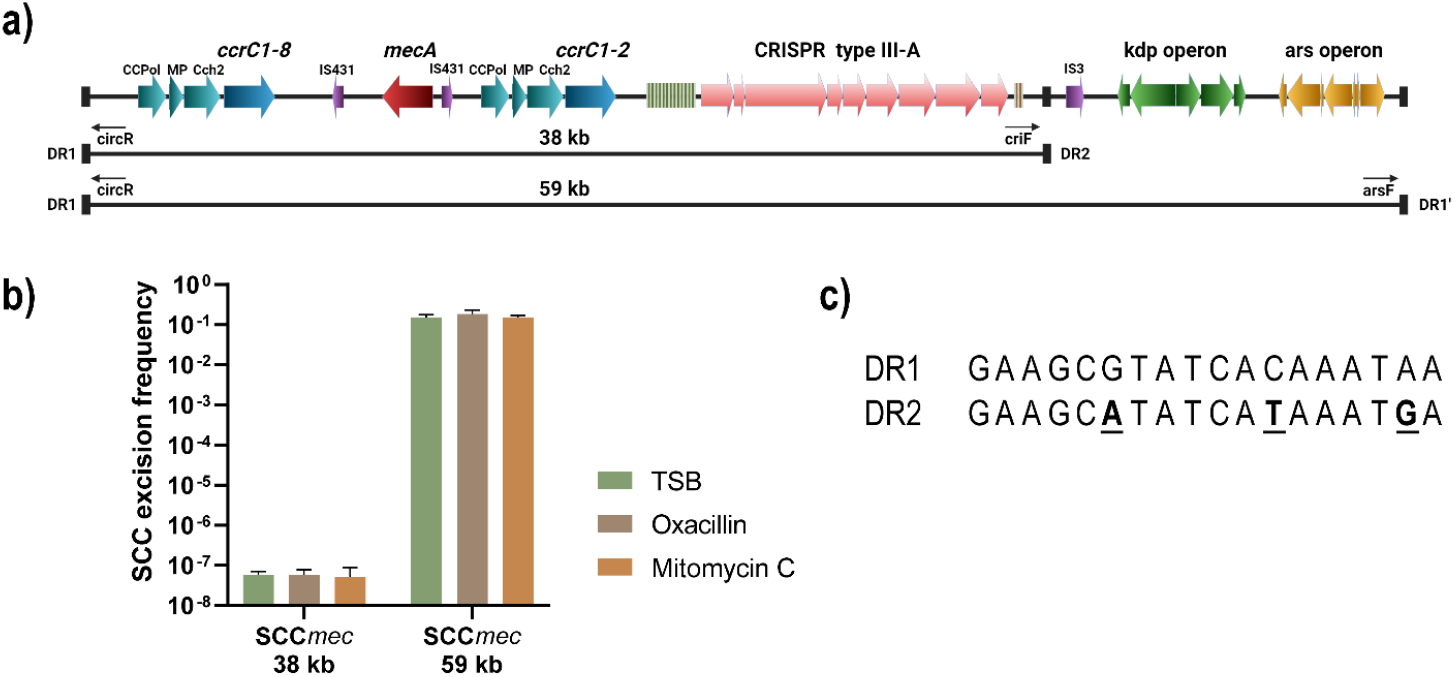
Excision of SCC*mec*. **(a)** The SCC*mec* type V(5C2&5) of 1109000 with predicted circularizable fragments. The primers and location of direct repeats (DR) are shown below the SCC*mec*. **(b)** Excision frequencies of the 38 and 59 kb SCC*mec* fragments compared to chromosomal DNA quantities (n=3). Primer pairs *criF/circR* and *arsF/circR* were used for detecting the 38 kb and 59 kb fragments, respectively as shown in (a), (c) Alignment of DR1 and DR2, with mismatches highlighted in bold and underlined in the DR2 sequence.

In *S. aureus*, horizontal gene transfer is often facilitated by phages [35], and we therefore tested if the circularized SCC*mec* could be transferred by phage transduction. To this end, we infected strain 110900 with the transducing phage φ11 and prepared phage lysates. We used the phage lysates to infect *S. aureus* strains RN4220, 8325-4 φ11, and Newman, and selected for transductants that had acquired the *mecA* gene and thereby would become oxacillin resistant. Despite repeated attempts we did not detect any transductants. Thus, under our experimental conditions, the SCC*mec* cassette does not transduce via phage φ11. This is likely caused by the limited packaging capacity of φ11, which may be unable to accommodate the 58 kb SCC*mec*. Furthermore, since the 38 kb fragment rarely circularizes, any phage-mediated transfer of this smaller fragment would probably be below the detection limit. Similarly, a recent study identified *S. aureus* transfer of an SCC*mec* cassette by natural transformation, however we were unable to show natural transformation of the SCC*mec* and CRISPR-*cas* in our strains [36].

### The ST630 Type III-A CRISPR-Cas system is active but inefficient against phage infection

To examine the anti-phage activity provided by the *S. aureus* type III-A CRISPR-Cas system, we deleted the CRISPR-*cas* locus in strain 110900 and infected wild type (WT) and ΔCRISPR mutant cells with the lytic phage phiIPLA-RODI that is targeted by spacer 1.9 in the CRISPR array (Table 2). At a multiplicity of infection (MOI) of 10^-6^ (1 phage to 10^6^ bacteria), strain 110900 survived the infection, whereas the ΔCRISPR mutant failed to grow (Figure 4). At higher initial phage concentrations (MOI of 10^-3^ and 10^-4^), the killing of the ΔCRISPR mutant correspondingly happened earlier than for the WT strain. Interestingly, for the WT 110900 there was a large variation between the technical replicates when cells were infected at MOI of 10^-5^ and 10^-6^, with some bacterial populations lysing from the phage infection, while others survived for longer periods of time. Thus, at low MOIs, the CRISPR-Cas system protects 110900 against phage killing, but at higher MOIs the system is overwhelmed.

**Figure 4.**
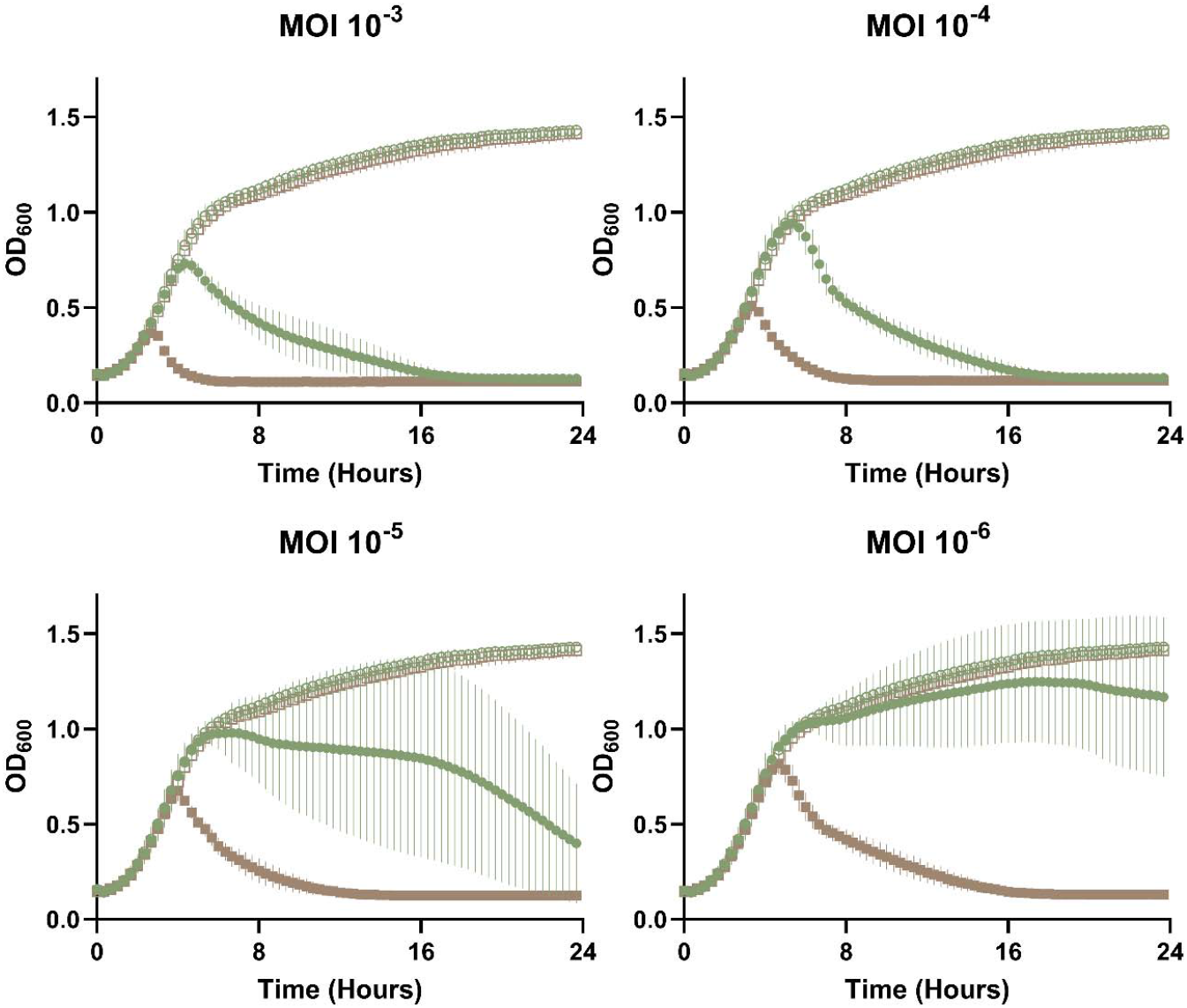
CRISPR-Cas activity against phage philPLA-RODI during liquid infection. Cultures (OD_600_ 0.15) of 110900 WT (green) or the ΔCRISPR mutant (brown) were grown with (closed symbols) or without (open symbols) increasing MOIs of philPLA-RODI phage in 1:1 TSB:phage buffer with OD_600_ being monitored every 20 min for 24 hours (n=3, 5 technical replicates).

Since we had observed high variability in CRISPR-Cas protection against phages at low MOIs, we investigated this further. Thus, we repeated the experiment shown in figure 4 by infecting 110900 with philPLA-RODI at an MOI of 10^-6^, and at the end of the experiment (24 h), sequenced the phages in the wells of the five technical replicates where lysis had occurred. In three of the replicates, we found that the phages had an identical 476 bp deletion covering AVU41_gp213, AVU41_gp212 and the intergenic region upstream of these two genes (see supplementary figure 1). Importantly, AVU41_gp213 is the philPLA-RODI gene that is targeted by the 110900 CRISPR spacer 1.9. Thus, escape mutants of philPLA-RODI can circumvent the CRISPR-Cas activity.

We also examined the burst size of philPLA-RODI when infecting either the WT 110900 or the ΔCRISPR mutant, using a one-step growth curve (Figure 5a). We found that the burst size of the phage was 17 when infecting the WT, but 33 in the ΔCRISPR mutant, showing that CRISPR-Cas reduces the average amount of phage progeny produced. Similarly, in a plaque assay when the two strains were used as recipients for a phage titer determination, significantly more plaque forming units (PFU) were detected on the ΔCRISPR mutant compared to the WT, equating to an efficiency of plaquing (EOP) of 29% on the WT recipient (Figure 5b). Thus, our data shows that the 110900 CRISPR-Cas type III-A system is active and offers protection against low MOI infection, however the system can be overwhelmed and can select for phage escape mutants where the targeted region has been deleted.

**Figure 5.**
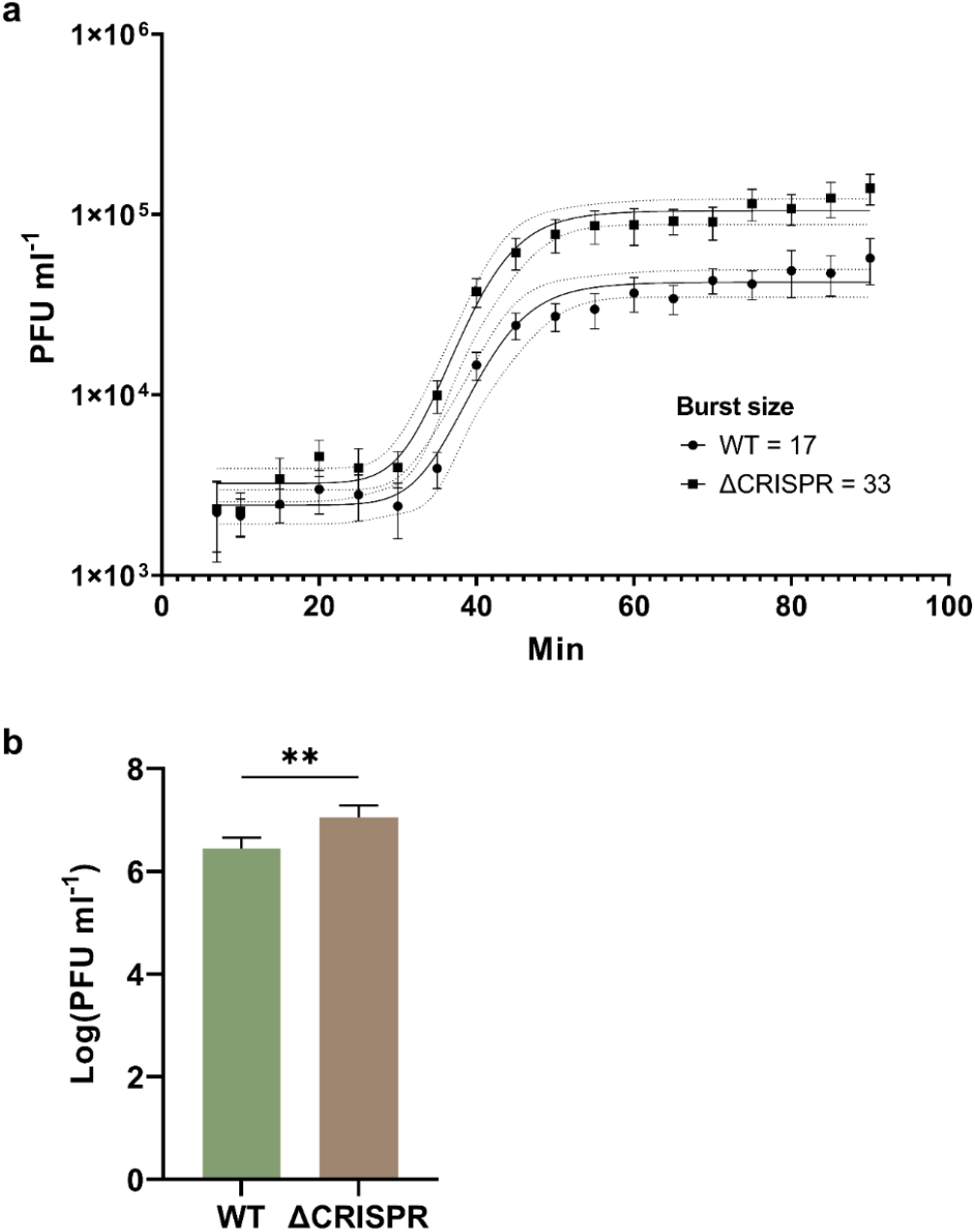
CRISPR-Cas reduces philPLA-RODI burst size. a) One step growth curves of philPLA-RODI using WT 110900 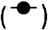 or the ΔCRISPR mutant 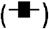 as propagative strain determined from 5 biological replicates (+/− SEM) and compared with a paired two-tailed t test (*p* value - <0.0001****). The nonlinear regression sigmoidal curves were fitted using GraphPad Prism with least squares regression. Burst size was calculated as the top plateau average divided by the bottom plateau average. b) Plaque formation of philPLA-RODI using either WT 110900 or the ΔCRISPR mutant as recipient shown as mean log transformed PFU ml^-1^ of 4 biological replicates +/− SD, the *p* value of the unpaired t test was 0.0087**. Efficiency of plaquing (EOP) of philPLA-RODI on 110900 WT was 29% of that on the ΔCRISPR mutant.

## Discussion

In *S. aureus*, CRISPR-*cas* systems are only found in a subset of strains. Here, we show that in Danish clinical MRSA isolates, 2.9% were positive for the CRISPR-*cas* type III-A system with the highest frequency in the emerging ST630 clone, where more than 50% of the isolates were CRISPR^+^ Inspection of the CRISPR arrays revealed conserved spacers at the 5’ end of the array and curiously, CRISPR array 1 mainly differed at the center of the array. This supports recent findings that the order of spacers in CRISPR arrays may arise from a combination of events, including middle-array insertion, recombination within or between arrays and horizontal transfer of all or part of the array [37]. As spacer adaptation via Cas1 and Cas2 has been observed only with overexpression in an inducible setting [38], the type III-A system could rely on alternative mechanisms to adapt new spacers such as recombination between CRISPR spacers and their cognate protospacer [39] or from integration and excision of temperate phages [40]. Also, low CRISPR adaptation frequencies in *S. aureus* may prevent self-targeting, and may permit horizontal acquisition of useful genes, such as phage-inducible chromosomal islands or plasmids. Indeed, *S. aureus* strains carry many prophages and PICIs on their genome that are important for virulence [41, 42].

We further observed that there is a high degree of conservation of spacers across staphylococcal species (Figure 2a) and between *S. aureus* strains (Figure 1). This has previously been observed for *S. aureus* strains [7] as well as between staphylococcal species [8, 43]. Besides suggesting a low frequency of adaptation events of the type III-A CRISPR-Cas system, the interspecies conservation of spacers as well as *cas* genes indicates horizontal transfer between staphylococci. As the type III-A CRISPR-*cas* system is located within similar SCC*mec* type V(5C2&5) cassettes across species, including *S. capitis, S. schleiferi* and *S. pseudintermedius*, this supports that the SCC*mec* cassette is of non-aureus origin [44]. Indeed, we observed that the *S. psuedointermedius* SCC*mec* containing CRISPR-*cas* has an identity of 99% to that of strain 110900. The occurrence of highly conserved SCC*mec* elements containing CRISPR-*cas* in strains of sequence type ST630 could be related to their unusual composition of cell wall teichoic acids that has been proposed to enable horizontal gene transfer between coagulase-negative staphylococci and *S. aureus* [21].

CRISPR-*cas* systems have previously been associated with mobile genetic elements including plasmids, genomic islands, and transposons [45, 46]. In *S. aureus*, the SCC*mec* cassette has been reported to excise from the genome [33] and we also observed that the entire SCC*mec* type V(5C2&5) including CRISPR-*cas* excised at a high frequency. Generally, it is unknown how SCC*mec* cassettes are transferred, but proposed routes include conjugative plasmids, transduction at low frequencies, or most recently via natural transformation [47]. We attempted to transduce the SCC*mec* type V(5C2&5) with the general transducing phage φ11 and transfer the element via natural competence, but in both cases, we were unable to detect transfer. Thus, it is unclear if and how SCC*mec* cassettes carrying CRISPR-*cas* are transferred between *S. aureus* strains.

Employing one of the strains identified in the screen, we analyzed the activity of the endogenous CRISPR-Cas system of the ST630 strain 110900 against phage philPLA-RODI, which is targeted by spacer 1.9 (figure 1 and table 2). The one step growth curves of the philPLA-RODI confirmed that CRISPR-Cas confers some protection against phage proliferation, with the philPLA-RODI having a burst size of 17 in the WT strain and 33 in the ΔCRISPR mutant. A previous study also indicated that type III-A CRISPR-Cas systems may influence burst size, with the temperate phage φNM1y6 yielding a burst size of ~5 PFU when propagated in a strain encoding a targeting spacer compared to ~85 PFU in the absence of targeting spacers [10]. Whereas the latter experiments were performed in *S. aureus* using a plasmid-encoded *S. epidermidis* CRISPR-Cas system with an engineered spacer targeting a virulent version of temperate *S. aureus* phage φNM1, our results show that native type III-A CRISPR-Cas systems also impact phage burst size.

In general, we found that the WT 110900 strain was more resistant to phage infection compared to the ΔCRISPR mutant both in liquid and plaque assays (Fig 4 and 5b, respectively). However, during infection of a liquid culture, the CRISPR-mediated protection was greatly dependent on MOIs, where the system was overwhelmed at MOIs of 10^-4^ and higher. At MOIs of 10^-5^ and 10^-6^, there was great variation in CRISPR-Cas-mediated protection, with some cultures surviving while others were killed by phage escape mutants that carried deletions of the 35 bp 1.9 protospacer sequence. This could be linked to the fact that the spacer 1.9 targets a gene in the long terminal repeats of the philPLA-RODI, which is presumably a late-expressed gene. For late-expressed genes, it has been shown that both the DNase and RNases of the type III-A system are required to prevent phage-mediated killing, and that many phage genomes can accumulate before targeted degradation occurs [10]. This could increase the likelihood of escape mutants occurring during phage replication. Escape from the type III-A system has been shown to stem from large deletions of the invading MGE that include the protospacer region [32]. It is curious that we saw identical regions deleted in 3 of our 5 phage escape mutants, and initially we thought that this deletion mutant of philPLA-RODI was present in the ancestral phage pool. However, we did not observe the escape mutations in our philPLA-RODI stock and indeed, with our experimental set-up, the likelihood of the mutations pre-existing in our phage inoculum is low, given that the number of phages added was approximately 7 PFU with the MOI of 10’^6^. An additional explanation for the conservation of the deletion mutants could be that this region is a hot spot for recombination. Despite incomplete CRISPR-Cas-mediated phage defense, the emergence of phage escape mutants demonstrates that the native CRISPR-Cas system does confer an evolutionary restraint for phages.

Interestingly, the level of protection provided by the endogenous CRISPR-Cas system was much less than reported in the literature where heterologous expression vectors are used. In a recent study of the *S. epidermidis* type III-A CRISPR system expressed from a plasmid in an *S. aureus* strain, there was a 10^3^ – 10^4^ PFU ml’^1^ reduction of phage abundance compared to the WT *S. aureus* strain without the plasmid [13]. Here, we find a CRISPR dependent reduction of ~10^1^ PFU ml’^1^, implying that this endogenous *S. aureus* CRISPR-Cas confers less protection. However, this is also lower than in a previous study where we examined the type III-A CRISPR-Cas system encoded by *S. aureus* ST630 strain TZ0912 [7]. Differences in CRISPR-*cas* expression, spacer placement and composition could underlie some of the difference in CRISPR-Cas targeting efficiency; the TZ0912 spacer 6 has 34/35 identical bp with the philPLA-RODI target gene, whilst the 110900 spacer 1.9 has 33/35.

Collectively, we find that the endogenous CRISPR-Cas type III-A system in *S. aureus* is active in protecting against phages but does so inefficiently. Furthermore, the conservation of spacers between *S. aureus* strains and even between staphylococcal species indicates that acquisition of new spacers is a rare event. When it does happen, however, it may involve recombination between invading mobile genetic elements and the excised copies of the SCC*mec* type V(5C2&5) [39]. Given that phages outnumber their bacterial hosts in natural habitats, such as in the human microbiome [48, 49], our results indicate that the immunity conferred by the type III-A CRISPR-Cas system in *S. aureus* only offers a partial protection from invading genetic elements such as phages. This may not be surprising as previously, these systems have been tested in isolation and under optimal conditions, whereas in a more natural set-up, CRISPR-Cas is likely one part of a multi-layered protection system. Indeed, the bioinformatic tools DefenseFinder and PADLOC show that the 110900 strain carries additional defense systems, including restriction modification, abortive infection, Serine/Threonine Kinase, and Dodola systems (Supplementary Table 1 [50–53]). Therefore, these systems are likely to defend against phage killing in a synergistic or additive manner not observed in heterologous systems [54]. Likewise, CRISPR-Cas may be differentially regulated depending on environmental conditions, e.g. activity could be enhanced in biofilms where bacteria may encounter phages in low numbers at which the CRISPR-Cas system is effective in protection. Overall, our findings indicate that the native *S. aureus* type III-A CRISPR-Cas system provides partial phage immunity and is likely part of the wider phage defense arsenal working in synergy. Moreover, our finding that the entire SCC*mec* type V(5C2&5) including CRISPR-*cas* excised at a high frequency suggests that methicillin resistance and CRISPR-Cas-mediated phage defense systems may be mobilized simultaneously under yet unrecognized conditions. This could prove a challenge for efficient use of phages for therapy of MRSA infections, particularly of the emerging clone ST630 in which we found 50% of the isolates to harbor CRISPR-*cas*.

## Materials and Methods

### Isolate collection and classification

All primary bacterial isolates are recovered from Danish patients who were admitted due to a healthcare-associated infection between 27^th^ of October 2017 and 12^th^ of March 2019. They were subjected to whole-genome sequencing and analyzed as described below. All isolates were typed at Statens Serum Institut as part of the national MRSA surveillance program. Isolates that contain *cas1* and *cas2* genes were selected for further exploration. The genome sequence of strain 110900 has been published [22].

### Whole-genome sequencing and analysis

All isolates were whole-genome sequenced on an Illumina MiSeq with 2 x 251 bp paired-end reads. Isolates were assembled using SKESA or SPAdes. CRISPRCasFinder was used to identify isolates that contain the *cas1* and *cas2* genes [24],

### Spacer analysis

Spacers from CRISPR-*cas* positive isolates were aligned using Clustal Omega and clustered by hierarchical clustering with Ward’s method based on 2 indel difference calculated by ape version 5.0. A maximum-likelihood phylogenetic tree was calculated using IQ-TREE with default settings running with bootstrap of 100 from SNPs called and filtered by NASP version 1.0. A dendrogram of 37 strains was constructed using the method with arithmetic means (UPGMA). Visualization of spacers was performed using plotly Graph Objects [55] and adjusted in Adobe Illustrator.

### Spacer sequences

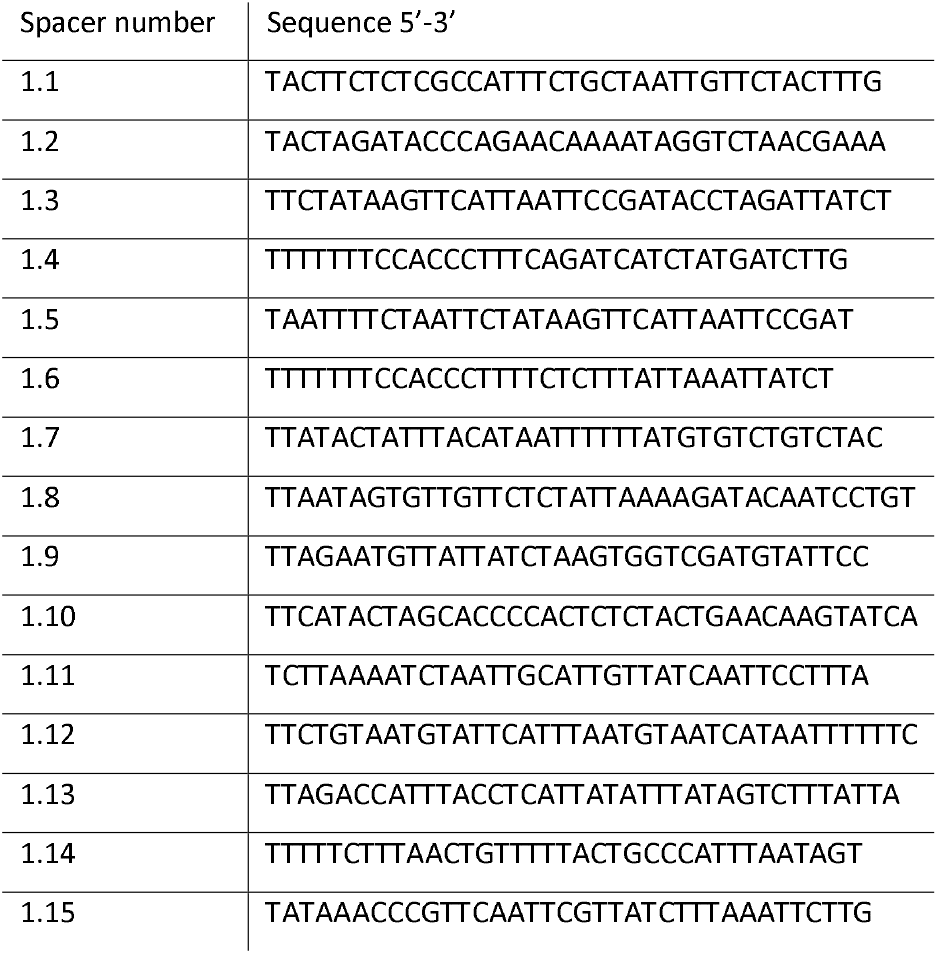
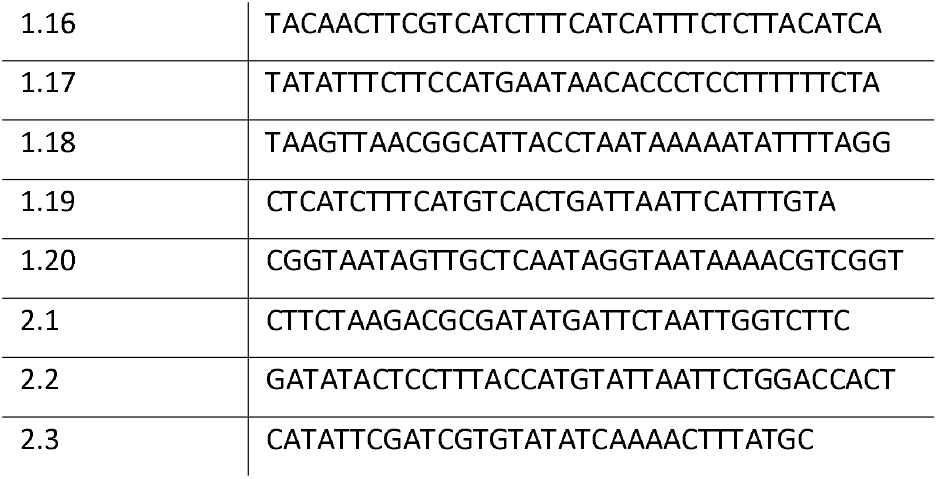

### Identification of spacer targets

The spacer groups were blasted against the entire NCBI database using the blastn algorithm on January 7, 2021. Bacterial isolates carrying the CRISPR-*cas* system and CRISPR arrays were not classified as protospacers. Hits representative of the targeted protospacer was included in Table 1.

### Chromosomal deletion of the CRISPR system

Sequences of 500 bp upstream and downstream of the CRISPR locus in strain 110900 were PCR amplified from purified 110900 chromosomal DNA with overlapping sequences. These fragments were used in a second overlap extension PCR reaction to construct a ~1 kb fragment, which was cloned into the PCR amplified pIMAY-Z plasmid backbone using a homemade seamless ligation cloning extract (SLiCE) as described by Monk and Stinear [56]. The mutant was created by following the step-by-step protocol also provided by Monk and Stinear [56].

### Liquid infection assay

Strain 110900 and the ΔCRISPR mutant were grown in TSB overnight and diluted to OD_600_ 0.15. The strains were then transferred into honeycomb bioscreen plates (95025BIO) in 125 μL aliquots and 125 μL of various phage philPLA-RODI lysate dilutions in phage buffer (1 mM MgSO_4_, 4 mM CaCl_2_, 50 mM Tris-HCl pH 8, 0.1 M NaCl), equivalent to MOIs 1 - 10^-7^, were added. The OD_600_ of each well was measured in a Bioscreen C instrument (Oy Growth Curves Ab Ltd) with measurements every 20 mins for 24 hours, at 30°C, with shaking. Shaking was paused 5 secs before each reading. For each experiment, 5 technical replicates of each condition were included and 3 biological replicates were performed in total. Positive bacterial growth controls of bacterial culture and phage buffer without phage (125 μL:125 μL), and negative controls of TSB and phage buffer (125 μL:125 μL) were included in each run. Results were plotted in an XY graph (GraphPad Prism 9) as mean +/− SD.

### Phage genome sequencing

DNA was extracted from a sample of the original WT philPLA-RODI phage lysate and 5 different philPLA-RODI samples grown on the CRISPR-positive 110900 strain for 24 hours in the bioscreen experiments, using the GenElute Bacterial Genomic DNA extraction kit (Sigma Aldrich), using the standard protocol with 200 μL phage lysate taken directly from the bioscreen plates. All isolates were whole-genome sequenced on an Illumina MiSeq with 2 x 251 bp paired-end reads. Sequences were assembled to the reference philPLA-RODI genome (NC_028765) using Geneious Prime with BBDUK trimmer plug in. SNPs and variations were called using Geneious Prime, excluding regions of high or low coverage, with any SNPs/variations present in the WT sample excluded from further analysis.

### SCC*mec* excision frequency assay

110900 was grown overnight and diluted to OD_600_ 0.05. Cultures were grown for two hours at 37 °C, where the cultures were either treated with antibiotics (oxacillin or mitomycin C, 0.5 μg/ml) or left untreated. After 1 hour additional incubation at 37 °C, 1 mL of culture was withdrawn and chromosomal DNA extracted using the Dneasy blood and tissue kit (Qiagen). The samples were normalized according to DNA concentrations and diluted 1:5, before being used in RT-PCR reactions. RT-PCR reactions were set up using the FastStart Essential DNA Green Master kit (Roche), using four different primer pairs; *criF/circR* (38 kb fragment circularization), *arsF/circR* (59 kb fragment circularization), *adsAF/adsAR* (chromosomal reference). Reactions were run on a LightCycler^®^ 96 instrument (Roche) and data was analyzed using the 2^-ΔΔCt^ method. The PCR product sequences were confirmed by cloning the PCR products into the pCR™4Blunt-TOPO^®^ vector by TOPO cloning (Thermo Fisher) and sequenced using the M13 reverse primer site by sanger sequencing (Eurofins).

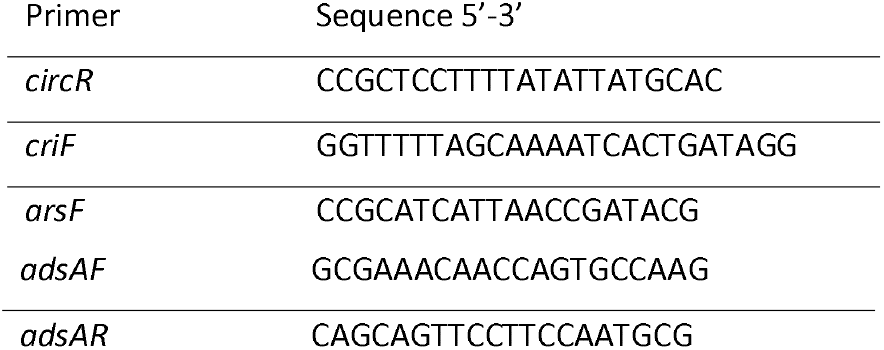

### SCC*mec* transduction experiment

ST630 isolate 110900 was grown overnight and diluted to OD_600_ 0.05. For initial phage infection of strain 110900, 200 mL cultures were grown at 37 °C until OD_600_ 0.15 and the cells were collected, before resuspension in 1:1 TSB:phage buffer, at a final volume of 100 mL. The cultures were infected with φ11 at various MOIs, at 30°C for 4 hours. If visual lysis of the culture was not complete, the culture was further incubated at room temperature overnight. Once visual lysis was complete, the lysates were filtered to remove any remaining bacterial cells (0.22 μm filters, Millipore stericup, PES).

Lysates were precipitated to increase the concentration of phage particles and any potential CRIPSR/SCC*mec* transductant particles. Lysates were incubated with DNase (2.5 U ml^-1^) and RNase (1 μg ml^-1^) at 37°C for 1 hour. NaCl was added (58.4 g ^-1^) and lysates incubated on ice with shaking for 1 hour. Lysates were centrifuged at 11,000 x g for 10 min at 4°C and the supernatant collected. PEG 8000 was added at 10% w/v and the lysates incubated on ice at 4°C overnight. The lysates were centrifuged at 11,000 x g for 10 min at 4°C and the supernatant discarded. Phage precipitants were resuspended in 1.6 mL phage buffer and quantified by titration methods.

Transduction assays were attempted using recipient strains RN4220, 8325-4 φ11, and Newman. Briefly, overnight cultures of recipients were diluted to OD_600_ 0.05 and grown to OD_600_ 1.4, before 1 mL recipient, 100 μL of phage lysate and 4.4 mM CaCl_2_ were incubated at 37°C for 20 mins. The mixture was plated out in 3 mL TSA top agar (50% agar) on TSA with oxacillin (0.4 μg mL^-1^) and sodium citrate (17 mM). Plates were incubated at 37°C for 24 hours and checked for colonies, if no colonies were observed the plates were incubated for an additional 24 hours and checked again.

#### One step growth curves

One step growth curves were performed essentially as previously described (Kropinski, 2018). Briefly, bacterial propagative strains were subcultured and grown to mid-log phase before adding 5 x 10^5^ PFU mL^-1^ philPLA-RODI to 9.9 mL culture for 5 minutes adsorption. Dilutions were performed as previously described and samples were taken every 5 minutes until 90 minutes. Plaque counts were normalized to the adsorption control and PFU mL^-1^ were calculated and plotted. Nonlinear regression curves were fitted to the data using the sigmoidal model in GraphPad Prism using the least squares method and the burst size was calculated by dividing the top plateau average by the bottom plateau average.

#### Phage titer assays

Phage titers were performed as previously described [57]. Briefly, recipient strains were grown to 0.35 OD_600_ and 100 μL aliquots of recipient mixed with 100 μL phage lysate, at different dilutions in phage buffer (10^0^ to 10^8^; 1 mM MgSO_4_, 4 mM CaCl_2_, 50 mM Tris-HCl pH 8, 0.1 M NaCl). After 10 min incubation at room temperature, 3 mL of liquid PTA (phage top agar; Oxoid nutrient broth no. 2, agar 3.5% wt/vol) was added and the mixture was poured out on PB plates (phage base; nutrient broth no. 2, agar 7% wt/vol). Plates were incubated at 37°C overnight and plaques were counted. EOP was calculated by dividing WT PFU ml^-1^ by Δ PFU ml^-1^ and multiplying by 100 to transform to percentages.

#### DefenseFinder and PADLOC bioinformatic analyses

The genome of 110900 (accession CP058615.1) was checked for the presence of defense systems using the defense system identification tools PADLOC [50, 51] and DefenseFinder [52], using the standard settings on their respective webservers. For PADLOC the input file was the NCBI-formatted genome (.gb), whilst for DefenseFinder the input file and pipeline was Nucleic fasta and analysis was preceded by Prodigal.

## Data availability

Sequencing data are available at doi: 10.5281/zenodo.7747092.

---

**Supplementary figure 1.**
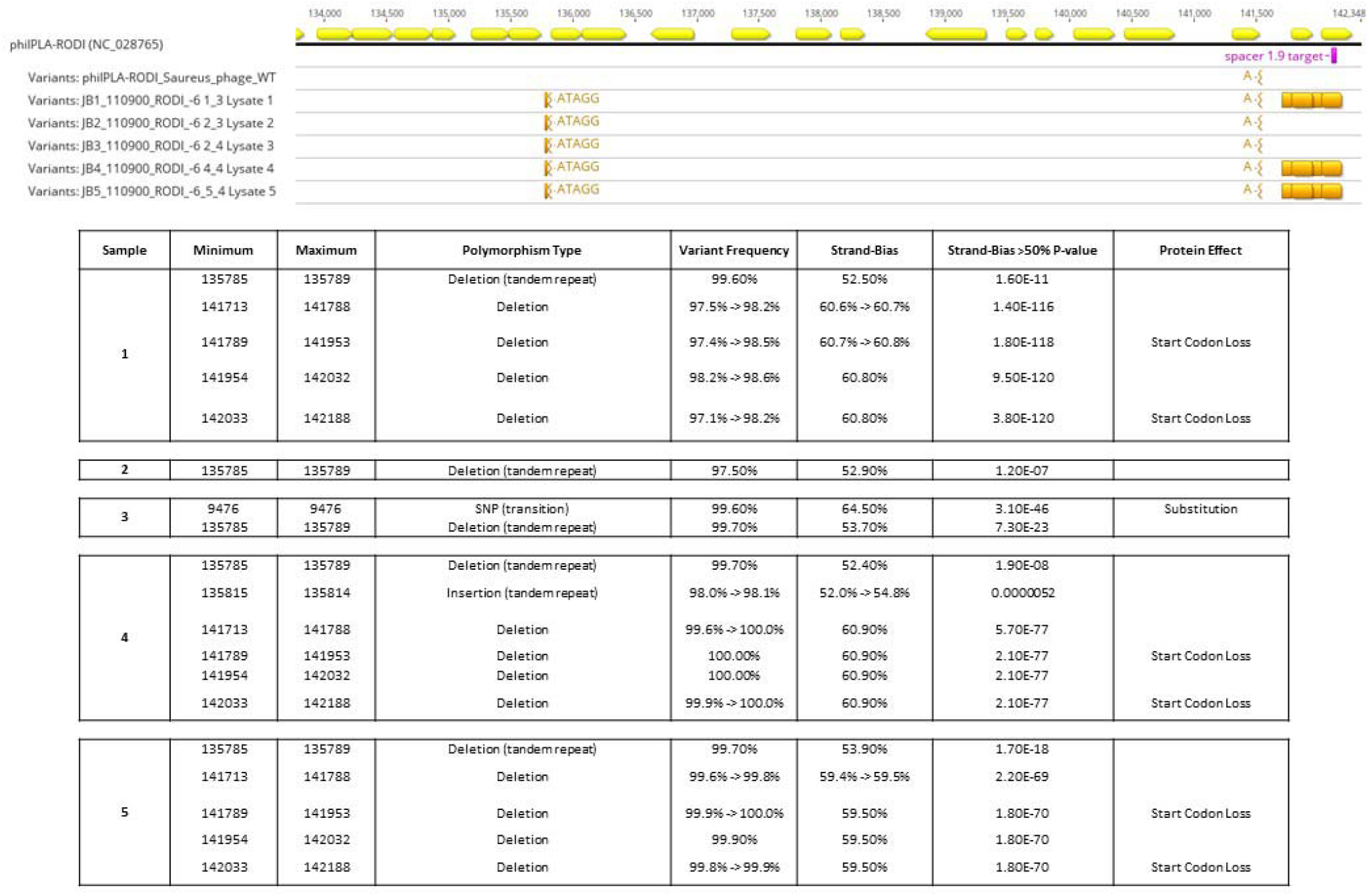
The majority of the philPLA-RODI lysates that escaped CRISPR targeting in the bioscreen assays contained deletions at the spacer targeting site. The schematic shows the alignment of the variants found following whole genome sequencing of our ancestral WT philPLA-RODI lysate and 5 separate samples of philPLA-RODI that lysed 110900 WT in the experiment shown in Fig 3. Spacer 1.9 targets the phage at the indicated protospacer sequence in the phage genome. The table below shows the variations in detail, excluding those also present in the sequenced WT philPLA-RODI or with an insignificant P-value. “Sample” indicates the philPLA-RODI sample number and minimum/maximum are the position in the phage genome.

**Supplementary table 2.** Results of PADLOC and DefenseFinder analyses of 110900 genome. Tables show the different potential defense systems identified in the genome of 110900 (CP058615.1) by the tools PADLOC [50, 51] and DefenseFinder [52]. System type is indicated by ‘Defense system’, ‘Protein name’ refers to the homologous defense protein, ‘Start’, ‘End’ and ‘Strand’ indicate position of the gene in the genome.

PADLOC
| Defense system | Protein name | Start | End | Strand |
| --- | --- | --- | --- | --- |
| cas_type_III-A | Cas1_IIIa | 62698 | 63604 | + |
|  | Cas2_IIIa | 63603 | 63909 |  |
|  | Cas10a | 63922 | 66196 |  |
|  | Csm2 | 66198 | 66624 |  |
|  | Csm3 | 66625 | 67270 |  |
|  | Csm4 | 67280 | 68189 |  |
|  | Csm5 | 68191 | 69214 |  |
|  | Csm6 | 69213 | 70482 |  |
|  | Cas6_IIIa | 70478 | 71213 |  |
| RM_type_IV | mREase_IV | 376053 | 378915 | + |
| RM_type_IIG | REase_MTase_IIG | 2019926 | 2021831 | - |
| DMS_other | Specificity_I | 2505372 | 2506584 | - |
|  | MTase_I | 2506576 | 2508133 |  |

| Defense system | Protein name | Start | End | Strand |
| --- | --- | --- | --- | --- |
| cas_type_III-A | Cas__cas1_I_II_III_IV_V_VI_5 | 62699 | 63604 | + |
|  | Cas__cas2_I_II_III_IV_V_VI_1 | 63604 | 63909 |  |
|  | Cas__cas10_III-A_1 | 63923 | 66196 |  |
|  | Cas__csm2gr11_III-A_1 | 66199 | 66624 |  |
|  | Cas__csm3gr7_III-A_1 | 66626 | 67270 |  |
|  | Cas__csm4gr5_III-A_3 | 67281 | 68189 |  |
|  | Cas__csm5gr7_III-A_4 | 68192 | 69214 |  |
|  | Cas__csm6_III_2 | 69214 | 70482 |  |
|  | Cas__cas6_I_II_III_IV_V_VI_7 | 70479 | 71213 |  |
| Stk2 | Stk2__Stk2 | 83716 | 84447 | + |
| Dodola | Dodola__DolA | 645903 | 646646 | + |
|  | Dodola__DolB | 646647 | 647783 |  |
| Abi2 | Abi2__Abi_2 | 2543259 | 2544197 | + |
| RM_type_IIG | RM_Type_IIG__Type_IIG | 2019927 | 2021831 | - |

## Acknowledgements

NMHK was supported by Lundbeck grant R264-2017-3936. HI and KM were supported by Danmarks Fri Forskningsfond (7017-00079B) and HI and JZB by Danmarks Fri Forskningsfond (0135-00271B). JTR was supported by EMBO Postdoctoral Fellowship ALTF 164-2021.

We thank Vi Phuong Thi Nguyen and Gitte Petersen for technical assistance in purifying *S. aureus* chromosomal DNA.

